# Development of Proton Density Fat Fraction Micro-MRI for Non-Invasive Quantitative Assessment of Bone Marrow Changes with Age and Radiation in Mouse Models

**DOI:** 10.1101/2025.01.03.631267

**Authors:** Hemendra Ghimire, Malakeh Malekzadeh, Ji Eun Lim, Srideshikan Sargur Madabushi, Marco Andrea Zampini, Angela Camacho, Weidong Hu, Guy Storme, Monzr M. Al Malki, Susanta K. Hui

## Abstract

**Background/Objectives:** Bone marrow (BM) adipocytes are critical in progressing solid tumor metastases and hematological malignancies across pediatric to aging populations. Single-point biopsies remain the gold standard for monitoring BM diseases, including hematologic malignancies, but are limited in capturing the full complexity of loco-regional and global BM microenvironments. Non-invasive imaging techniques like Magnetic Resonance Imaging (MRI), could offer valuable alternatives for real-time evaluation of BM diseases in both preclinical translational and clinical studies.

**Methods:** We developed a preclinical proton density fat fraction (PDFF) MRI technique for quantitative BM composition assessment, focusing on fat fraction (FF) within mouse femurs. We validated this method using aging mice and young mice subjected to 10 Gy X-ray irradiation, compared with young unirradiated mice as controls. Water-fat phantoms (0% to 100% fat content) were used to optimize the imaging sequence, and immunohistochemical (IHC) staining with H&E validates equivalent adipose content in the femur BM regions.

**Results:** Significant differences in FF were observed across age groups (p = 0.001 for histology and p = 0.0002 for PDFF) and between irradiated and control mice (p = 0.005 for histology and p = 0.002 for PDFF). A strong correlation (R^2^ ∼ 0.84) between FF values from PDFF and histology validates the accuracy of the technique.

**Conclusions:** These findings demonstrate the potential of PDFF MRI as a non-invasive real-time imaging biomarker for quantifying BM fat fraction in preclinical mice model studies, particularly in evaluating the effects of aging, disease progression, and irradiation therapy in pediatric and translational oncology research.

## 1 Introduction

Bone marrow (BM) adipocytes create a unique microenvironment that significantly influences the progression of solid tumor metastases (e.g., breast and prostate cancers) and hematological malignancies (e.g., leukemia and multiple myeloma) ^1-5^. These adipocytes support tumor growth and survival by supplying lipids and secreting adipokines, thereby influencing the BM niche to promote malignancy. Aging-related increases in marrow adiposity further exacerbate these effects^6^. Current BM monitoring relies on invasive techniques like single-point biopsies or aspirations, which are limited in capturing the spatial heterogeneity and complexity of the BM microenvironment^6-8^. These invasive methods also fail to provide dynamic changes in marrow composition and function during disease progression or therapy in both preclinical and clinical settings. Therefore, developing non-invasive imaging techniques in preclinical mouse models is essential for providing high-resolution, comprehensive insights into the spatial and functional dynamics of BM. Such advances would enable a more comprehensive understanding of adipocyte interactions, cancer progression, and therapeutic responses, ultimately supporting precision diagnostics, and improved treatment strategies.

Magnetic Resonance Imaging (MRI), along with other technologies like computed tomography (CT)^9^, dual-energy X-ray absorptiometry (DXA)^10^, and optical imaging, have enabled in vivo quantification of BM composition^11-13^. MRI is particularly valuable for adipose composition analysis because it provides high-resolution images without exposing patients to ionizing radiation. Techniques like PDFF MRI measures BM fat by precisely distinguishing between fat and water signals using chemical shift encoding^14-16^. Clinical studies have increasingly focused on the applications of PDFF-MRI for BM composition and heterogeneity studies. However, its feasibility and accuracy in preclinical translational studies remain underexplored, particularly as mice have small BM space, and lower BM fat compared to humans^17^, presenting unique challenges in spatial resolution.

Achieving accurate PDFF measurements at higher magnetic field strengths, such as 7 Tesla (7 T), is complex and requires multiple gradient echoes with short echo-to-echo spacing to accurately capture the relative in-phase and out-of-phase signals of water and fat. Similar to the development of other MRI imaging techniques, such as T2-MRI to assess BM inflammation^18^, Dynamic Contrast-Enhanced MRI (DCE-MRI) to examine leukemia-induced changes^19^, and multiparametric approach to quantify myeloproliferative neoplasms associated^20, 21^, PDFF-MRI application of mice femur BM requires further validation. Establishing standardized protocols and correlating PDFF-MRI findings with histological data is crucial for determining its utility for early-stage studies and bridging the gap between preclinical research and clinical applications.

This article presents the development of PDFF-MRI for assessing BM adiposity in mouse femurs using aging mice, young mice exposed to 10 Gy X-ray irradiation, and control groups. Mouse models provide valuable insights into BM adiposity^17^, as studies indicate that it increases with age^22^ and X-ray irradiation influences marrow adipocyte repopulation^11^. The study follows a three-step process: (1) evaluating PDFF with a water-fat phantom linearity test; (2) conducting PDFF-MRI of femur BM in live mice; and (3) validating findings through histological quantification of adipose in the local area to achieve the purpose of the study: to evaluate the feasibility of the PDFF-MRI technique for mouse femur BM evaluation. Herein, a phantom study is essential for optimizing imaging settings and ensuring accurate PDFF measurements, which minimizes potential errors in in vivo studies^23^. Furthermore, correlating PDFF-MRI results with histological data is essential for confirming the technique’s reliability in preclinical research^17, 24^. This validation approach is essential for optimizing imaging protocols, advancing hardware and translational methodologies, and improving our understanding of disease mechanisms and treatment responses in mouse models before transitioning to human trials.

## 2. Materials and Methods

The study employed a cross-sectional design to investigate the feasibility and efficacy of monitoring fat fraction in mouse femurs using PDFF-MRI imaging. All studies were performed in accordance with the Institutional Animal Care and Use Committee at City of Hope (COH) National Medical Center Duarte, CA. Mouse models representing aging, X-ray irradiation, and corresponding young unirradiated controls were used in the study.

### 2.1. Phantom Preparation

Water-fat phantoms with varying fat percentages (by volume): 0%, 5%, 10%, 20%, 40%, 60%, and 100%, were prepared using mixtures of gelatin, deionized water, emulsifier washing liquid, and commercially available peanut oil. Peanut oil was chosen due to its proton nuclear magnetic resonance similarity to triglycerides in adipose tissue^25^. A solution of 0.1596 grams of 225 bloom gelatin (Specialty food source, Brockton, MA) per milliliter of deionized water was autoclaved at 121°C for 15 minutes, then cooled to 50 – 55°C and constantly fixed within this temperature range on the laboratory hot plate magnetic stirrer with magnetic mixer for further steps^26, 27^. The gelatin and variable amounts of peanut oil were vortexed to emulsify the mixture to create homogeneous solutions. Then we used the Benchmark Scientific BV1000 Bench Mixer Vortex Mixer (115V) at its maximum speed of 3200 RPM for varying durations based on concentration: 5 minutes for a 5% fat concentration and up to 60 minutes for a 60% fat concentration. To remove microbubbles, we used an ultrasonic bath (Branson Ultrasonic Corporation CPXH). We used Micro CT images (Precision X-ray) with a pixel size of 0.1×0.1 mm^2^ to check the microbubbles.

### 2.2. Mouse Models

For aging effect study, 93 weeks aged C57BL/6J (JAX 000664) mice were purchased from Jackson Laboratory (Maine, USA) and housed at the COH animal facility. By the time of MRI imaging, these mice were 96 weeks old.

For X-ray irradiation effect study, 9-12-week-old C57BL/6N (NCI-Charles River Laboratory) mice were used. The mice were divided into two groups: no treatment control (5 mice), and right leg irradiated (5 mice), underwent right leg irradiation with 10 Gy X-ray using a 3D image-guided focused X-ray irradiation system (X-RAD SmART Precision X-ray machine, Precision X-Ray, North Branford, CT, USA), operating at 225 kV and 13 mA.

### 2.3. Mouse Preparation and Monitoring for MRI Scanning

Mice underwent anesthesia with 2-4% isoflurane at a flow rate of 2 L/min of oxygen throughout the imaging. Mice were kept warm at 37°C by the Minerve multi-station temperature control unit (Esternay, France) throughout the experiment. The respiration rate was maintained at 40-50 /minute and monitored by an MR-compatible small animal monitoring and gating system (SA Instruments, Inc., NY, USA) through a respiration pad taped on the mouse. Initially, imaging development used a surface coil positioned over the femur region for improved signal. However, the surface coil setup was time-consuming and required careful temperature management to prevent drops in body temperature, which is challenging over longer sessions. The body coil with built-in temperature control allowed for stable physiological conditions control, reducing stress and avoiding hypothermia. Therefore, a mouse body coil was employed with regular mouse bedding. To ensure proper femur alignment in a supine position, flat popsicle stick slabs were utilized to bridge the two femurs in the same plane, ensuring coplanarity.

### 2.4. Image Acquisition and Analysis

A multi-echo gradient echo sequence was acquired with 6 different echo times, field of view (FOV) = 22.55×30.00 mm^2^, slice thickness = 1 mm, spacing between slices = 0.10 mm, matrix size = 128×128, time repetition (TR) = 180 ms, echo times (TEs) = 3760, 4080, 4400, 4720, 5040, 5360 µs, flip angle = 25°, bandwidth = 66.7 KHz, number of excitations (NEX) = 2, and pixel size= 0.1758×0.2344 mm^2^. A water-fat decomposition of the images was performed via the preclinical scan software in MR Solutions scanner, the gradient-based optimization for organ-specific estimation (GOOSE) algorithm. This algorithm accurately separates fat and water signals for FF quantification. The main feature of this algorithm is the ability to reduce, swaps artifacts in fat water decomposition problems^28^.

For FF analysis, we used MATLAB R2024b to create region-specific masks on T2-weighted images. These masks were precisely co-registered to match the corresponding locations and orientations in both the histological sections and PDFF-MRI images, ensuring accurate comparison and analysis of the same anatomical area.

### 2.5. Tissue Preparation

Following MRI imaging, femurs were harvested from euthanized mice and processed for histological analysis. Freshly dissected mouse femurs were fixed in 10% paraformaldehyde (PFA) overnight (24 hours) at 4°C (to preserve the tissue components and morphology).

### 2.6. Hematoxylin & Eosin (H&E) Staining and Histological Analysis

The femurs were washed three times with phosphate-buffered saline (PBS) and then decalcified in 10% ethylenediaminetetraacetic acid (EDTA) solution for 48 hours at room temperature. After decalcification, the femurs were rinsed with PBS and embedded in paraffin to create tissue blocks. Tissue slides were deparaffinized, rehydrated, and stained with Modified Mayer’s H&E Y Stain (America MasterTech Scientific) on an H&E Auto Stainer (Prisma Plus Auto Stainer, SAKURA) according to standard laboratory procedures. H&E-stained slides were scanned using a NanoZoomer S360 Digital Slide Scanner (Hamamatsu) and whole slide images were viewed by NDP.view 2 image viewer software. Subsequently, Fiji/ImageJ was employed to generate representative images. MATLAB R2024b was used to quantify the area occupied by adipocytes within the region as covered by mask created to analyze FF of PDFF MRI images within the BM section. Distal femurs were chosen for validation due to their ease of handling, especially in the brittle femurs of aged or irradiated mice, and for their relatively higher adipose content, allowing for precise sectioning and analysis.

### 2.7. Statistical Analysis

Statistical analyses were performed using GraphPad Prism (GraphPad Software Inc., La Jolla, CA, USA). Group comparisons were conducted with one-way ANOVA and Tukey’s post hoc test, with significance set at p < 0.05. R^2^ values were calculated in Excel to evaluate the relationship between histology and PDFF-MRI data via linear regression as well as phantom study. Data are expressed as mean ± SEM, and significance levels were indicated as: ns = not significant, ^*^p < 0.05, ^**^p < 0.01, ^***^p < 0.001.

## 3. Results

### 3.1. Technological Eevelopment for MRI-based Assessment of FF in Mouse Femur BM

The technological development of MRI techniques for assessing FF in mouse femur BM is shown in Figure 1. This development enabled accurate, artifact-free PDFF MRI imaging, which was crucial for preclinical translational studies focused on BM adipose content. Figure 1 (A) presents the schema of the experimental design, showing the steps and timing for handling the mice and acquiring images. Since proper alignment of the femur’s central core is key for accurate BM adipose content imaging, both prone and supine positions were tested (Figure 1.B). Supine positioning was more effective for easier femurs alignment for complete BM visualization, and this positioning is followed for MRI study. To reduce motion artifacts from respiration, a custom bedding platform to use with a surface coil was initially developed (Figure 1.D). However, a standard bedding setup used for body coil was ultimately preferred because it made adjusting the field of view (FOV) easier without affecting image quality. The axial and coplanar alignment of the femurs was validated through CT scans followed by MRI, ensuring accurate positioning of femurs for precise BM imaging.

**Figure 1.**
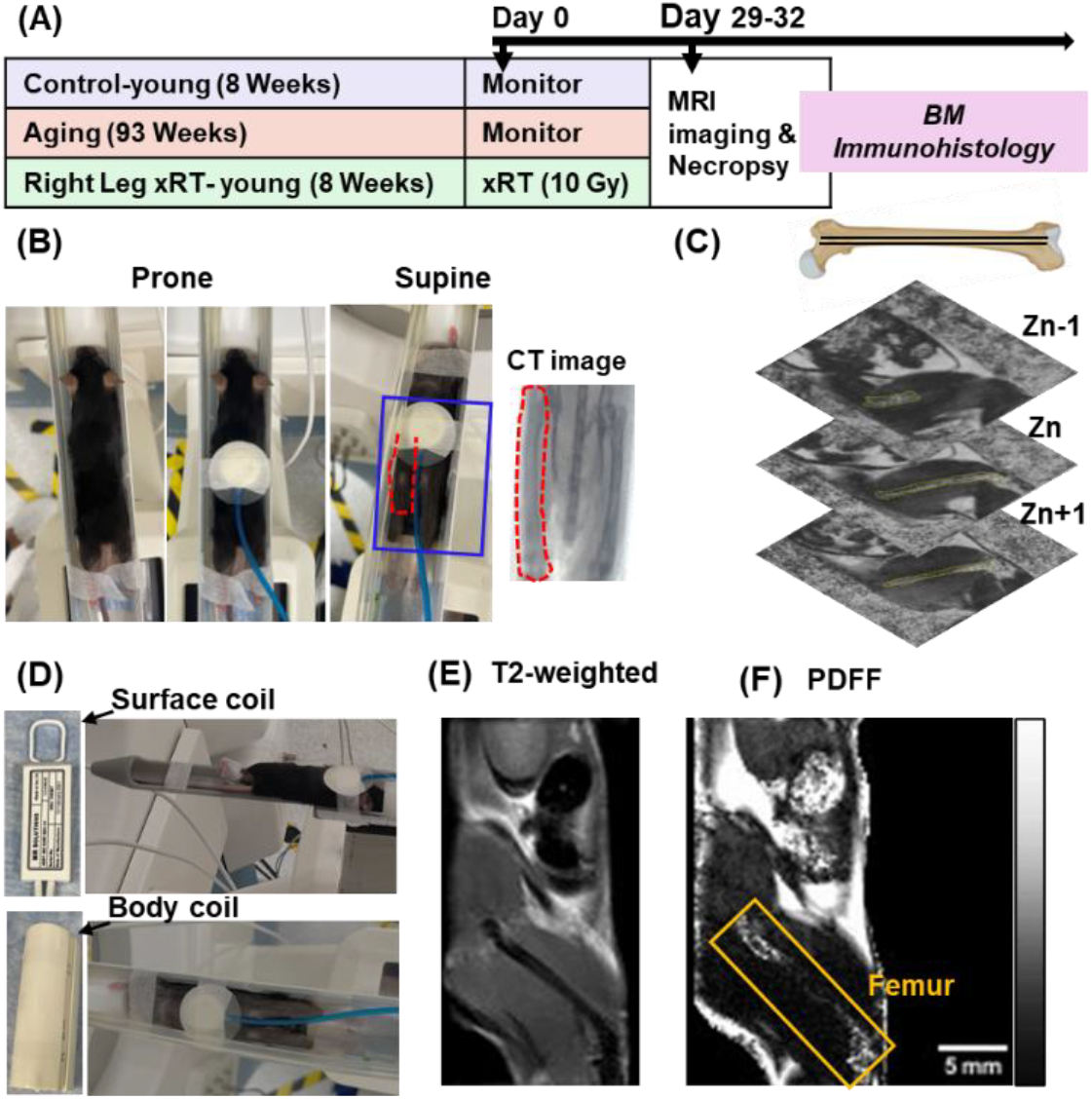
Technological development for MRI-based assessment of FF in mouse femur BM. (A) Schema of the experimental design, showing the handling of mouse and tissue samples over time. (B) The mouse is positioned in both prone and supine orientations for optimal axial alignment of femurs and marrow within the imaging plane. The supine positioning allows for the easier connection to ECG gating, and precise temperature control. The accurate positioning of the femurs for entire region of marrow imaging is further confirmed through CT scans, followed by MRI imaging. (C) Optimization of imaging parameters, including resolution, slice thickness, and inter-slice spacing to have a clearer visualization of fat content within BM region. Z-stack slicing in the aligned femur was illustrated using femur prepared in BioRender. (D) Comparative testing of custom-built surface (femur) coils and a conventional body coil for PDFF MRI. The body coil was selected due to its similar sensitivity for PDFF-MRI imaging and the additional benefit of maintaining animal warmth. (E-F) Representative T2 weighted (E), and PDFF (F) images showing the right femur BM in sagittal orientation. Herein, representative T2 weighted and PDFF images were prepared using a 19-week-old healthy mouse.

Optimization of MRI parameters, including resolution, slice thickness, and inter-slice spacing, was vital for obtaining clear images of the small adipose regions within the BM. Parameters were set up such that at least one z-stack slice covers the marrow region of entire distal to proximal femur (Figure 1.C). The FOV was specifically focused on the lower portion of the mouse’s body, including the right femur, to ensure comprehensive imaging of the marrow region. Figures 1 (E) and 1 (F) provide representative sagittal images, including T2-weighted (Figure 1.E) and PDFF (Figure 1.F) scans. These images demonstrate the effectiveness of the imaging protocol in producing detailed, artifact-free PDFF images suitable for further analysis.

### 3.2. Optimization of the MRI Imaging Sequence with Phantom

The water-fat phantom studies demonstrated strong correlations between the known fat fraction and the measured PDFF values obtained from a 7 T preclinical MRI system, with an R^2^ value of 0.9831 (Figure 2. A). There is a slight overestimation in pure water sample while using 7 T preclinical MRI system, aligning with findings reported in the literature^29^, which suggest that similar discrepancies can occur in fat quantification techniques.

**Figure 2.**
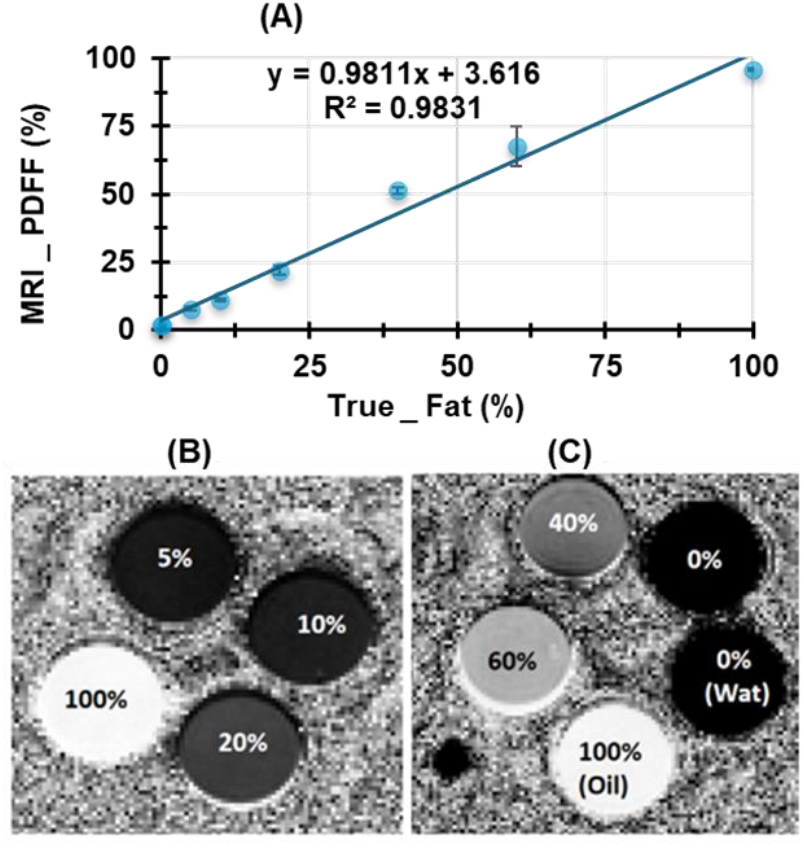
Water-fat phantom study showing strong correlation between measured and true FF. (A) Correlation between true fat volume fraction and the measured mean FF using MRI. (B) FF maps display fractions of 5%, 10%, 20%, and 100% fats. (C) FF maps indicating 0% (water), 0% (Gelatin & Water), 40%, 60%, and 100% (oil).

The FF maps for fat concentrations of 5%, 10%, 20%, and 100% effectively demonstrate the system’s differentiation capabilities (Figure 2.B). Additionally, Figure 2 (C) illustrates the imaging response to varying compositions: 0% (water), 0% (gelatin and water), 40%, 60%, and 100% (oil). The standard deviation observed in the 60% fat sample is likely to be due to microbubbles (<100 μm) influencing the measured values. These results confirm the efficacy of the 7 T preclinical MRI system in assessing fat fractions in phantom models.

### 3.3. Representatives T2-weighted and PDFF MRI Sequences from Each Study Groups

Figure 3 shows the T2-weighted, PDFF, and merged PDFF/T2 MRI sequences to highlight BM characteristics within the right femur region of different mouse models. The merged PDFF/T2 images provide a comprehensive view of the bone and surrounding tissues, making it easier to identify FF distribution within the femur BM. Figure 3 (A) shows representative images from a healthy young control mouse showing a mid-plane view of the right femur through T2 weighted, PDFF, and merged sequences. These images serve as the baseline where BM exhibits normal, relatively low levels of fat, reflecting an optimal hematopoietic environment. Figure 3(B) presents images from mice with irradiated legs, where noticeable changes in FF are observed, particularly in the distal femur region. This FF increase suggests irradiation-induced disruption of the marrow’s cellular environment, as fat accumulation is often associated with impaired hematopoiesis and reduced regenerative capacity. Such changes may reflect a shift toward a less supportive marrow niche, which could impact recovery and immune function post-radiation^30^. Similarly, aging leads to a significant rise in fat content within BM, particularly in the distal femur (Figure 3.C). Age-related fat accumulation within BM changes the marrow’s microenvironment, often leading to a decline in its hematopoietic potential and shows age-associated bone conditions^22^.

**Figure 3.**
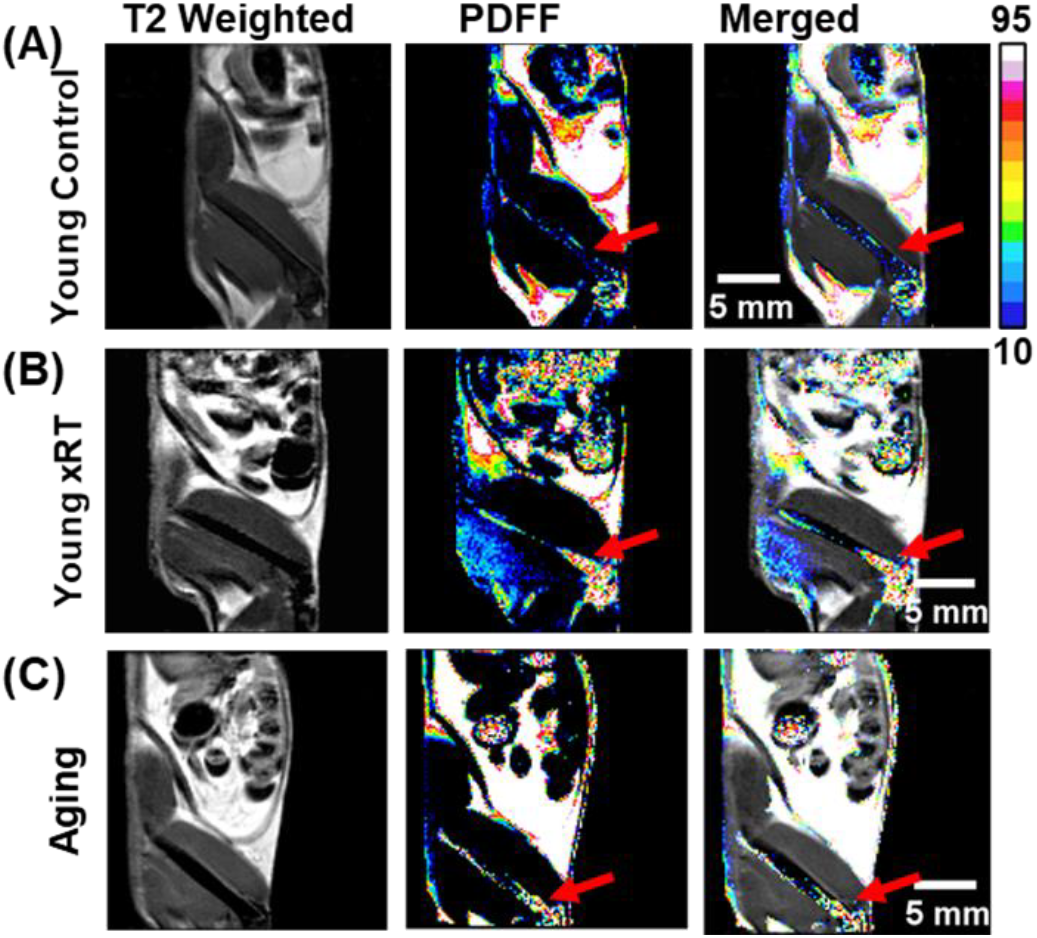
T2-weighted and PDFF MRI sequences of the mouse body, focusing on the BM region in the right femur. (A) Representative images from a young control mouse, showing a mid-plane view of the right femur in T2 weighted, PDFF, and merged images. (B) Images of an irradiated mouse model, showing irradiation-associated changes in FF within the corresponding femur BM. (C) Images from an aged mouse, demonstrating age-related changes in fat distribution. The PDFF and merged images provide enhanced visualization of fat distribution within the body and femur BM due to both irradiation and age. The red arrows point to the distal femur marrow, where FF levels are consistently higher compared to mid-shaft and proximal areas in both irradiated and aged mice.

### 3.4. Technological Validation with Distal Femur BM Histology

Figure 4 shows the quantitative analysis of BM adiposity in H&E-stained distal femur sections and its correlation with PDFF-MRI findings from the same regions. Figure 4 (A) shows a representative PDFF image of an aging mouse highlighting with differential fat content, where abdominal fat (red square) shows ∼90% FF, and femur muscle (blue square) shows <12% FF. This image reflects the anatomical variability in fat content. Figure 4 (B) shows mean FF from the ROI in the femur BM regions of each group (n=5 per group), derived by carefully selecting regions of interest (ROI) from T2-weighted images. Aging mice exhibit the highest FF in femur BM, highlighting a significant age-related accumulation of fat cells within the marrow. This finding aligns with evidence that increased BM adiposity, especially in aged or irradiated models, may impair hematopoiesis and contribute to reduced marrow resilience. The differences in BM fat content among aging, irradiated, and control groups were statistically significant (P < 0.0013).

**Figure 4.**
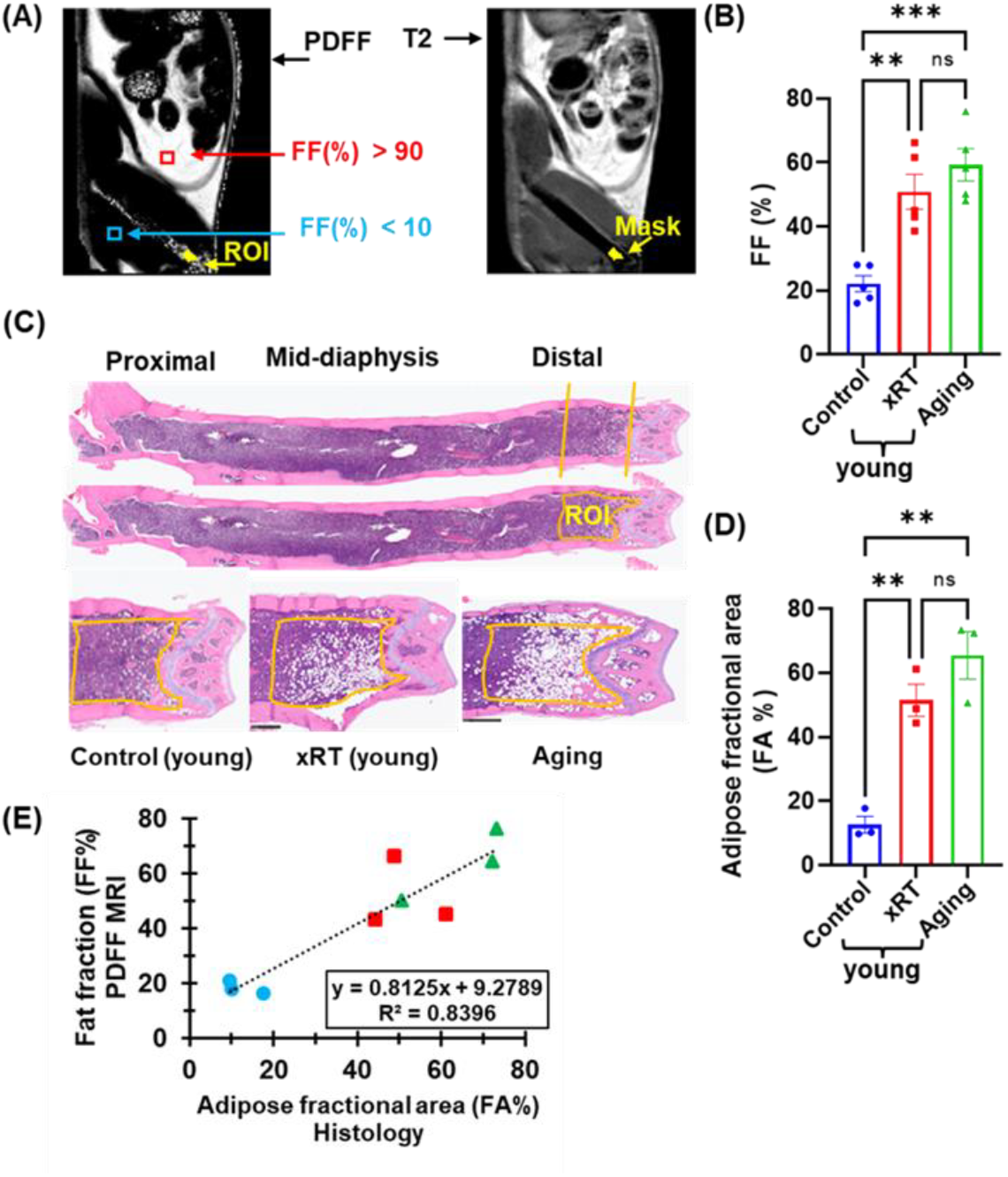
Quantification of FF (%) in distal femur BM of studied mouse groups. (A) Representative PDFF image of the right femur side of an aging mouse. Abdominal fat exhibits higher than 90% FF (highlighted by the red-lined square), while femur muscle (blue square) is less than 12%. A mask was created using a T2 weighted image, focusing on a 1.5-2 mm region within the marrow, starting 1 mm away from the distal femur (lateral condyle). (B) Quantitative FF (%) in the region of interest (ROI) of young control (Control), young irradiated (xRD), and aging mice using PDFF MRI images. FF (%) significantly increases in ROI of irradiated and aging mice compared to control mice (P < 0.0013). (C) H&E-stained femur BM sections (proximal, mid-diaphysis, and distal parts) from a control mouse. The H&E-stained distal section (yellow-lined region) represents a 1.5-2 mm area near the popliteal surface, adjacent to the synovial membrane reflection line, which corresponds to the same region used for FF quantification in PDFF MRI imaging. Comparisons were made between young control, 10 Gy X-ray irradiated, and aging mice. (D) Quantification of the adipose fractional area in the distal femur BM (highlighted in (C)), showing significantly increased adiposity in irradiated and aging mice compared to controls (P < 0.01). Three mice (n = 3) were used in each group. (E) Correlation between the adipose fractional area from H&E-stained images and the integrated density from PDFF-MRI images. A strong correlation is observed, with an R^2^ value of 0.84, indicating consistency between histological and MRI-based fat quantification. Data are shown as mean ± SEM, and significance levels were indicated as: ns = not significant, ^*^p < 0.05, ^**^p < 0.01, ^***^p < 0.001.

H&E-stained histological sections of mouse femurs were further evaluated for BM adiposity calculation. The distal femur which is comparatively easier for sectioning without damaging local bone marrow for histology and is the position where adipocytes are highly populated (highlighted region) is used for further analysis as shown in Figure 4(C). Same as in PDFF images, the adipocyte population is more in the femur BM of aging and irradiated leg of young mice, while control mice have less. Figure 4(C) shows the representative H&E-stained BM region images of the distal femur, with highlighting region of analysis, where the fractional area covered by adipose is obvious in magnified black and white sections. Adipose fractional area (FA) is further quantified for each studied group to show the statistical significance of adipose alteration.

The femur BM in aging mice legs and irradiated legs of young mice have significantly high (p < 0.01) adipose composition compared to control (Figure 4.D). Figure 4(E) demonstrates a strong correlation (R^2^ = 0.84) between adipose fractional area from H&E-stained sections and FF% derived from PDFF MRI, validating the accuracy of MRI for assessing BM fat. This correlation confirms the consistency between histological and MRI-based fat measurements, with an R^2^ value indicating that over 84% of the variance in histological data is explained by the MRI results, supporting the use of PDFF MRI for BM fat quantification in preclinical studies.

## 4. Discussion

The feasibility of preclinical MRI for assessing PDFF within the femoral marrow is demonstrated by monitoring local FF in aging, young and irradiated young mouse models. A water-fat phantom was initially used to optimize key imaging parameters such as resolution and number of averages, for the GOOSE algorithm. The phantom study is essential for this validation, as it provides a controlled environment to adjust imaging settings and ensure the accuracy and reliability of PDFF measurements, minimizing potential errors when transitioning to *in vivo* applications^23^. Furthermore, the technical feasibility of PDFF-MRI of femur BM was validated through histological images of local regions. PDFF-MRI measurements of marrow FF strongly correlated with adipose volumes observed in histology of local region (R^2^ ∼ 0.84). These findings suggest the potential of PDFF-MRI to provide precise, real-time insights into marrow composition, particularly its adipose tissue, for understanding the BM pathophysiology. Importantly, the appropriate selection of mouse models of diseases is critical for assessing biological variability, an advantage that cannot be performed with synthetic phantom studies^31^. The present study includes phantom and diverse preclinical mouse models relevant to pediatric oncology to capture adipose variability in BM region.

The heterogenous nature of the BM microenvironment, especially its adipose tissue and niche, is recognized as a sanctuary for disease progression and has attracted interest in conditions like hematologic malignancies, osteoporosis, sickle cell disease (SCD), and marrow failure syndromes^32^. Our previous studies showed that leukemia progression and residual disease influence BM niche components, including local FF[4], and the BM composition changes with age^12, 13^. These findings suggest that temporal and spatial variations in marrow FF could serve as potential imaging biomarkers for BM-related health issues, aiding in early detection, disease monitoring, and treatment planning^30^. Additionally, FF alteration due to therapeutic interventions, including conditioning regimens for hematopoietic stem cell transplantation (HCT), needs thorough translational validation. Advanced radiation modalities like total marrow irradiation (TMI)^33^, and total marrow and total lymphoid irradiation (TMLI)^34^ are being evaluated for their potential to control disease in the BM through dose-escalation while limiting dose to the vital organs like lung, liver, gut requires preclinical validation^35^. Evaluating the effects of therapeutic radiation on bone and marrow is crucial for developing precision treatments that minimize BM damage^30, 36-38^. Reverse translational from clinical to preclinical settings is often necessary for deeper insights into the complexities of treatments and diseases^31^. Therefore, developing reliable PDFF-MRI techniques for mouse models is critical to capturing the regional and global complexities of BM, including adipocyte distribution, hematopoietic cellularity, and tissue functionality. These non-invasive imaging approaches overcome the limitations of traditional biopsies, enabling earlier detection, improved disease monitoring, and more effective therapeutic interventions for BM-related conditions, which will advancing translational pediatric oncology.

Our developed technique allows for simultaneous adipose assessment of femurs and muscle fat within a single scan, making it ideal for longitudinal studies. Despite the promising results, there are several limitations in this study warrant discussion. First, while PDFF-MRI offers a non-invasive approach to monitoring marrow FF, its spatial resolution is lower compared to histological techniques, limiting its ability to capture finer molecular or cellular-level changes. To address this limitation, future studies could focus on using high resolution MRI and improving MRI protocols by optimizing acquisition parameters or employing advanced machine (deep) learning techniques for analysis. Second, the acquisition of data with excellent signal-to-noise ratio is a prerogative for accurate FF estimation, as noise-induced oscillations can otherwise lead to swaps in the fat water decomposition. Thus, further validation is needed to analyze the robustness of PDFF-MRI against noise levels. Additionally, our study primarily focuses on cross-sectional analysis, and the dynamics of BM fat content during disease progression or treatment are not fully explored. Longitudinal studies with more frequent imaging sessions would be crucial to better understand how marrow composition evolves over time in response to therapies.

## 5. Conclusions

The application of micro-MRI for assessing PDFF within the femoral marrow is demonstrated by using mouse models of aging, radiation treatments and young controls. The use of non-invasive PDFF-MRI in preclinical mouse model studies is a major step forward to addressing the critical limitations of traditional invasive biopsy analysis to understand complex interplay of marrow adiposity, disease progression, and treatment effects. Future research should focus on refining this imaging technique, exploring its longitudinal applications, and validating its findings in parallel with human studies. With its ability to provide real-time, quantitative insights into BM fat, PDFF-MRI has the potential to transform how we study BM diseases into translational oncology research, and beyond.

## Author Contributions

Conceptualization, S.K.H.; methodology, H.G, M.M., M.A.Z, A.C., J.E.L., S.S.M. and W.H.; software, H.G. and M.M.; validation, H.G., M.M. and J.E.L.; formal analysis, H.G. and M.M.; investigation, S.K.H. and S.S.M; writing—original draft preparation, H.G.; writing—review and editing, H.G., M.M., M.A.Z, W.H., G.S., and S.K.H.; visualization, H.G. and S.K.H.; supervision, S.K.H. and M.M.A.M.; project administration, S.K.H; funding acquisition, S.K.H. All authors have read and agreed to the published version of the manuscript.

## Institutional Review Board Statement

The animal study protocol was approved by the Institutional Animal Care and Use Committee (IACUC) of City of Hope (COH) National Medical Center Duarte, CA, USA.

## Data Availability Statement

The datasets used and/or analyzed in this study are available from the corresponding author upon reasonable request.

## Acknowledgments

The authors would like to acknowledge the valuable contributions of the Small Animal Imaging Core (SAIC) and Radiation Research Service Core (RRSC) at City of Hope National Medical Center.

